# Age-dependent changes in the dynamic functional organization of the brain at rest – a cross-cultural replication approach

**DOI:** 10.1101/2022.08.20.504632

**Authors:** Xi Yang, Xinqi Zhou, Fei Xin, Benjamin Becker, David Linden, Dennis Hernaus

**Affiliations:** Department of Psychiatry & Neuropsychology, Maastricht University, Minderbroedersberg 4-6, 6211 LK Maastricht, The Netherlands; Institute of Brain and Psychological Sciences, Sichuan Normal University, Chengdu, Sichuan, China; School of Psychology, Shenzhen University, Shenzhen, Guangdong, China; The Center of Psychosomatic Medicine, Sichuan Provincial Center for Mental Health, Sichuan Provincial People’s Hospital, University of Electronic Science and Technology of China, Chengdu, China

**Keywords:** Aging, Dynamic functional connectivity, Graph theory, Sliding window analysis

## Abstract

Age-associated changes in brain function play an important role in the development of neurodegenerative diseases. Although previous work has examined age-related changes in static functional connectivity (FC), accumulating evidence suggests that advancing age is especially associated with alterations in the dynamic interactions and transitions between different brain states, which hitherto has received less attention. Moreover, conclusions of previous studies in this domain are limited by suboptimal replicability of resting state fMRI and culturally homogenous cohorts. Here, we investigate the robustness of age-associated changes in dynamic functional connectivity (dFC) by capitalizing on the availability of fMRI cohorts from two cultures (Western European and Chinese). In both cohorts we consistently identify two distinct connectivity states: a more frequent segregated within-network connectivity state (state I) and a less frequent integrated between-network connectivity state (state II). In both cohorts, older (55-80 years) compared to younger participants (20-35 years) exhibited lower occurrence of and spent less time in state I. Moreover, older participants tended to exhibit more transitions between networks and greater variance in global efficiency. Overall, our cross-cultural replication of age-associated changes in key dFC metrics implies that advancing age is robustly associated with a reorganization of dynamic brain activation that favors the use of less functionally-specific networks.

**Highlights:** Aging is associated with a reorganization of dynamic functional brain connectivity.

Age-dependent dynamic functional connectivity changes are relatively stable across cultures.

Dynamic properties are promising neural indexes for brain aging in older healthy populations.

## 1. Introduction

As the world population ages, more and more individuals suffer from age-related neurodegenerative diseases, such as Alzheimer’s and Parkinson’s disease. Understanding how the brain changes with advancing age may therefore provide important insights into neurobiological processes underlying these diseases. Numerous studies have employed fMRI (functional magnetic resonance imaging) to examine age-related structural and functional brain changes, as well as their associations with cognitive decline (Betzel et al., 2014; Luo et al., 2020).

With respect to functional organization, functional connectivity (FC) - a commonly-used technique that computes pairwise correlations in the blood oxygen level dependent (BOLD) signal between voxels or large-scale networks - has been widely applied to examine age-related changes. Studies in this domain have reported an age-associated reduction in connectivity strength within large-scale networks of the brain, affecting among others the default mode (DMN), frontoparietal control (FPN), salience (SN), and executive control (CEN) networks (Betzel et al., 2014; Geerligs et al., 2015; La et al., 2015; Luo et al., 2020; Mak et al., 2017; Ng et al., 2016; Oschmann and Gawryluk, 2020). These findings are paralleled by a relative increase in between-network FC, for example between DMN and FPN, or DAN (dorsal attention network) and subcortical networks (Das et al., 2021; Ferreira et al., 2016; Geerligs et al., 2015; Stumme et al., 2020; Zonneveld et al., 2019). Results from these and other studies have led to the hypothesis that brain aging may be associated with decreased segregation of functionally-distinct networks, thus reflecting a reduction in the specialization of brain networks (Das et al., 2021; Ferreira et al., 2016; Grady et al., 2016; Stumme et al., 2020). Exacerbations of these patterns have been reported in individuals that suffer from Alzheimer’s and Parkinson’s disease, suggesting that neurodegenerative processes may accelerate the reduction in functional specialization of brain networks typically seen with normal brain aging (Cai et al., 2020; Cassady et al., 2021).

These previous studies primarily focused on static (i.e., time invariant) FC that examines spatial correlations over the entire MRI data-acquisition of several minutes, whereas they neglect temporal fluctuations in the intrinsic organization of the brain. Increasing evidence indicates that neural activity at rest is highly dynamic. That is, voxel-wise correlations of BOLD connectivity fluctuate over the course of an MRI session, taking on distinct configurations that vary from more sparsely-connected connectivity states that anatomically overlap with well-defined networks (e.g., CEN, DNM) to highly-interconnected states that resemble interactions between these networks (Allen et al., 2014; Preti et al., 2017). Dynamic functional connectivity (dFC) enables the investigation of these time-variant FC patterns and has been increasingly employed in healthy subjects and individuals that suffer from neurologic and psychiatric conditions (Allen et al., 2014; Li et al., 2019; Long et al., 2020; Zhang et al., 2021a).

Accumulating evidence suggests that aging is accompanied by changes in dFC patterns (Tian et al., 2018; Viviano et al., 2017; Xia et al., 2019; Zhang et al., 2021b). Xia et al. found that older individuals exhibited reduced variability in dynamic brain activation, as reflected in decreased switching rates between distinct connectivity patterns (Xia et al., 2019). Moreover, Tian et al. showed that older individuals spent more time in FC states characterized by weaker within-network connectivity - interpreted as less efficient information transfer - including less time spent in FC states dominated by within-network connectivity of sensorimotor and CEN (Tian et al., 2018). These studies suggest that age-associated reductions in the differentiation of static FC networks may be the result of more fundamental changes in the dynamic reorganization of and interactions between these networks.

Although these initial studies suggest that dFC is a promising technique that allows insights into mechanisms associated with age-dependent changes in brain function, it is unknown whether its findings are robust to a number of important challenges and confounds. It is increasingly recognized that resting state fMRI has low replicability (Bossier et al., 2020; Zuo et al., 2019). Whereas replicability issues may be partly attributed to the unconstrained nature of the resting state paradigm and analytical choices (e.g., sliding window length, head motion parameters, vascular signals) (Allen et al., 2014; Preti et al., 2017), the predominant focus on culturally homogenous samples from Western (European) cultures also strongly impedes the generalizability of previous findings.

Exposure to environmental risk- and resilience factors for age-related cognitive decline is culturally-dependent and may account at least partly for the lack of consistent results in this domain across studies (Bittner et al., 2019; Ewers et al., 2021). Important examples include cultural differences in dietary habits over the lifetime and health care accessibility for the elderly (Anderson et al., 2011; Govindaraju et al., 2018; Papanicolas et al., 2021). Ideally, we still hope to find cross-cultural commonalities in brain aging in these differentiated contexts. As a developing country, China differs from European countries in dietary habits, incomes, and healthcare systems. The determination of aging-related dFC that are invariant to these cultural/social differences would represent promising candidates for generalizable biomarkers (Fang et al., 2020; Zhang et al., 2015). Recent increases in the number of publicly-available fMRI data repositories enable the conduction of cross-cultural replication studies, thus allowing us to test the generalizability and replicability of age-related dFC changes across more diverse cultural contexts.

In the current study we assessed the cross-cultural robustness of dFC to identify age-related changes in dynamic brain connectivity. We analyzed resting state fMRI data from two independent and comparably large samples in Europe and China (LEMON: Leipzig Study for Mind-Body-Emotion Interaction Dataset - Germany and SALD: Southwest University Adult Lifespan Dataset - China) to obtain commonly-used dFC outcome measures after strictly controlling for methodological confounds. Our specific goals were to investigate the robustness of age-related changes in 1) temporal properties of dFC (fraction time, mean dwell time, and the number of transitions between distinct networks), and, 2) dynamic topological organization of functionally-connected networks (i.e. global and local efficiency) (Kim et al., 2017; Xia et al., 2019).

## 2. Materials and Methods

### 2.1 Participants

#### 2.1.1 Primary cohort

LEMON: Leipzig Study for Mind-Body-Emotion Interaction (Babayan et al., 2019) (http://fcon_1000.projects.nitrc.org/indi/retro/MPI_LEMON.html) This study comprised 228 healthy native German adults. Given the sensitivity of dFC to head motion (Preti et al., 2017), we employed stringent head motion criteria (max. head motion> 3 mm translational, > 3° rotation; n=3). We additionally excluded participants I) that were taking psychoactive drugs (urine drug screening and semi-structured interview for, buprenorphine, amphetamine, benzodiazepine, cocaine, methamphetamine, morphine/heroine, methadone, and marihuana; n=19), 2) with neurological disorders (DSM-5 diagnosis; n=28) and/or alcohol abuse and family history of alcohol addiction (DSM-5 diagnosis and semi-structured interview; n=53), 3) with failed spatial normalization as identified by visual inspection (n=2), and, 4) with missing demographic and/or MRI data (n=8). After applying these criteria a total of 133 participants remained, in line with previous studies (Doucet et al., 2021), both dataset subjects were subdivided into two distinct participant groups (young: n=88, age-range = 20-35, male/female=57/31; older: n=45, age-range=55-80, male/female = 22/23). All participants provided written informed consent. The study was approved by the ethics committee of the medical faculty at the University of Leipzig, Germany.

#### 2.1.2 Cross-cultural replication cohort

To conduct a cross-cultural replication of our results we used the publicly-accessible SALD resting state fMRI dataset (Wei et al., 2018) (Southwest University Adult Lifespan Project, (http://fcon_1000.projects.nitrc.org/indi/retro/sald.html). Out of the available 494 datasets of healthy native Chinese adults, one hundred sixty-three participants were excluded based on head motion (max. head motion > 3 mm translation, > 3° rotation; n=16), failed spatial normalization as defined by visual inspection (n=21), ages that fell outside of the age ranges of the young and older LEMON participants (n=127) and missing fMRI data (n=1). Please note that some participants fit multiple exclusion criteria, the numbers reported here therefore reflect the number of participants that met each exclusion criteria. Three hundred thirty subjects were included for further data analysis (young: n=168, age-range=20-35, male/female=68/100; older: n=162, age-range=55-80, male/female=66/96). All participants provided written informed consent. The study was approved by the Research Ethics Committee of the Brain Center of Southwest University (see Table 1).

**Table 1 legend:**
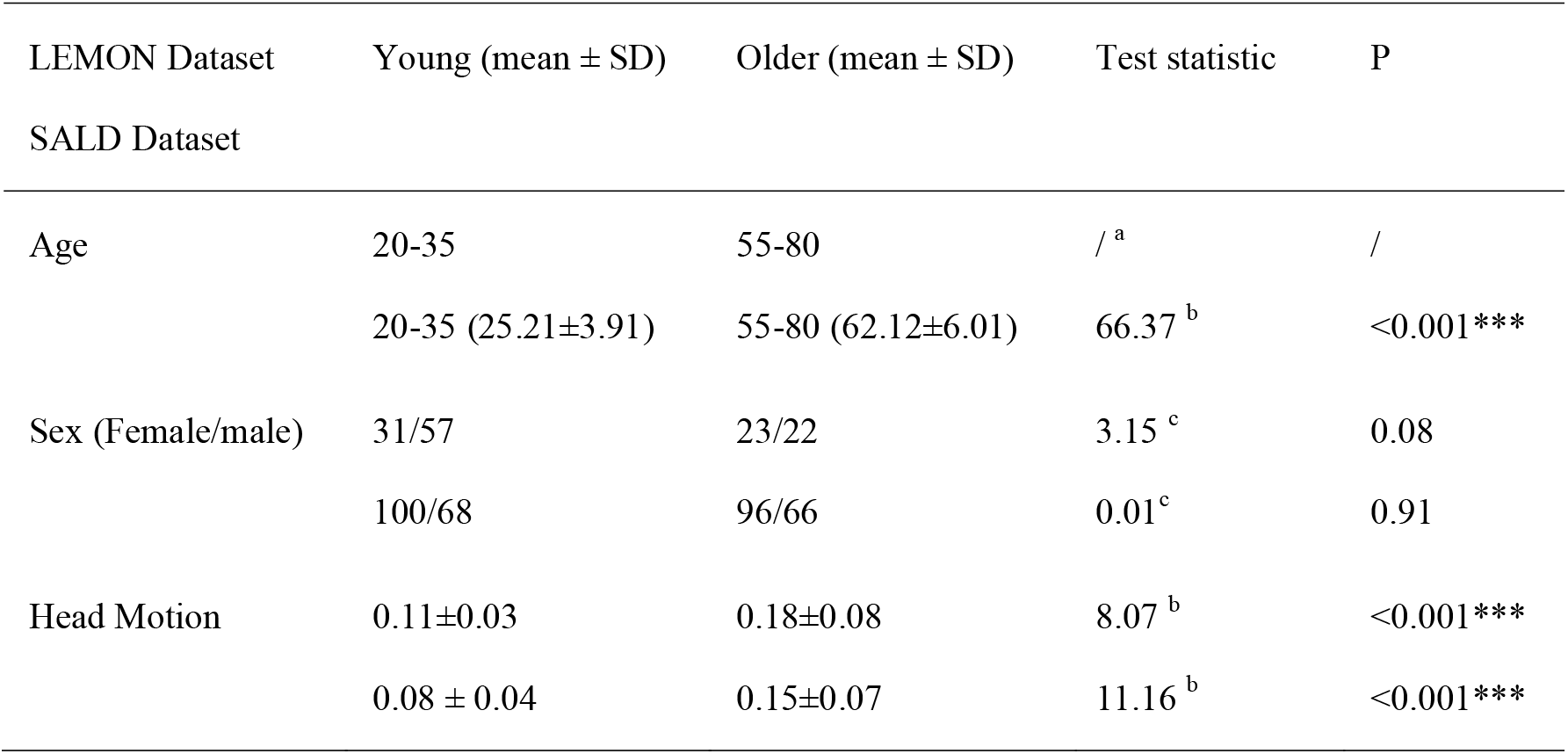
Descriptive statistics for both datasets. a: the LEMON dataset only contained age ranges; b: two samples T-test; c: chi-square test. ***: p< 0.001.

### 2.2 fMRI acquisition

#### 2.2.1 Primary cohort

MRI data were collected on a 3 Tesla MRI system (MAGNETOM Verio, Siemens Healthcare GmbH, Erlangen, Germany) equipped with a 32-channel head coil. Structural T1-weighted images were acquired using a Magnetization-Prepared 2 Rapid Acquisition Gradient Echoes (MP2RAGE) (Marques et al., 2010) sequence (TR=5000 ms, TE=2.92 ms, FA 1=4°, FA 2=5°, FOV=256 mm, voxel size=1×1×1 mm). Resting state data were acquired using a T2*-weighted Echo Planar Imaging (EPI) sequence with the following parameters: TR=1400 ms, TE=30 ms, FA=69°, FOV=202 mm, voxel size =2.3×2.3×2.3 mm, slice thickness=2.3 mm, image matrix=88×88. 15 min 30 s (657 volumes) of resting state data were collected for each participant. During the scan, participants were asked to keep their eyes open while looking at a low-contrast fixation cross.

#### 2.2.2 Cross-cultural replication sample (validation cohort)

All SALD MRI data were collected on a 3 Tesla MRI system (Siemens Magnetom Trio, Siemens Medical, Erlangen, Germany) equipped with a 16-channel whole-brain coil. Structural data were acquired using a magnetization-prepared rapid gradient echo (MPRAGE) sequence (TR = 1900 ms, TE =2.52 ms, FA =9 °, FOV =176, and voxel size = 1×1×1 mm). FMRI data were acquired using a gradient echo-planar image (GRE-EPI) sequence (TR = 2000 ms, TE = 30 ms, FA = 90°, FOV=220 mm, voxel size = 3.4 × 3.4 ×4 mm, slice thickness = 3 mm, image matrix = 64×64). The total acquisition time was 8 min 4 s (242 volumes). Participants were instructed to remain awake with their eyes closed and to rest without thinking of anything.

### 2.3 fMRI data analysis

#### 2.3.1 MRI data preprocessing

All resting state fMRI data were preprocessed in a combined SPM12 (http://www.fil.ion.ac.uk/spm/; Wellcome Trust Centre for Neuroimaging) and DPABI (Version: V4.3_200401; http://rfmri.org/dpabi) (Yan et al., 2016) analysis pipeline. We discarded the first five volumes to ensure MRI signal equilibrium. Since scan duration can influence resting state connectivity estimates (Birn et al., 2013), we discarded the last 312 volumes of resting state data for participants in the LEMON dataset, resulting in a scan duration (7 min 56 s, 340 volumes) that was nearly identical to the scan duration of the SALD dataset (dFC results using the entire timeseries are reported under Supplementary Results 1). Next, all images were corrected for head motion, spatial normalization using the Diffeomorphic Anatomical Registration Through Exponentiated Lie Algebra (DARTEL) template, after which the resulting images were smoothed (6 mm full-width at half-maximum Gaussian kernel [FWHM]). We discuss band-pass filtering as part of the dFC procedures reported below (see 2.3.3).

Comparing head motion as defined by framewise displacements (FD) (Jenkinson et al., 2002), we observed a significant difference between the young and older participant groups in the LEMON (young: mean ± SD = 0.11±0.03; older: mean ± SD = 0.18±0.08; p<0.001, two samples T-test) and SALD dataset (young: mean ± SD= 0.08 ± 0.04; older: mean ± SD = 0.15 ± 0.07; p < 0.001, two samples T-test). We therefore included 24 motion parameters (6 motion parameters of the current volume and the preceding volume, plus each of these values squared) for each participant. In addition, nuisance signals were calculated using CompCor (component-based noise correction method), resulting in five principal components from white matter (WM) and cerebrospinal fluid (CSF) (Behzadi et al., 2007).

#### 2.3.2 Group Independent Component Analysis

Next, group-level ICA was performed using the Group ICA Of fMRI Toolbox (GIFT) (http://icatb.sourceforge.net/) (Calhoun et al., 2001) with the aim of identifying networks of interest. For subject-specific data reduction, a total of 120 principal components were retained using principal component analysis (PCA). For the group level data reduction, the concatenated (reduced) subject-level data were decomposed into 100 group independent components (ICs) using the expectation maximization algorithm. The Infomax ICA algorithm was repeated 20 times in ICASSO (http://www.cis.hut.fi/projects/ica/icasso) to improve stability (Himberg et al., 2004). Components with an average intra-cluster similarity value >0.8 were retained (Kim et al., 2017). To obtain subject-specific spatial maps and time courses for each independent component, group independent components were back-projected using the GICA back reconstruction algorithm (Calhoun et al., 2001).

Of the 100 independent components, 51 and 43 ICs - for the LEMON and SALD dataset, respectively - were selected according to Allen’s criteria (Allen et al., 2014) based on consensus between three independent evaluators (X.Y., X. Z., F. X.). Components were identified based on the principles that 1) IC peak coordinates of spatial maps were primarily in gray matter and showed low spatial overlap with vascular, ventricular, motion, and other artifacts, 2) time courses were dominated by low-frequency fluctuations (ratio power <0.10 Hz to 0.15–0.25 Hz), and, 3) time courses were characterized by a high dynamic range (dynamic range >0.02). Based on those criteria, we additionally calculated spatial correlations between independent components and template to re-identify significant components (correlation coefficient >0.2) (Du et al., 2020).

#### 2.3.3 Computation of dFC

DFC was estimated using the GIFT toolbox via two steps: 1) a sliding window approach (i.e., changes in FC over time) and, 2) K-means clustering (i.e., recurrence of FC activity patterns). Resting state time series were divided into bins of 40 TRs (i.e.,40*1.4=56 s) with sigma 3-TR of Gaussian, and a 1 TR step between windows. Additional processing steps were applied to improve signal-to-noise ratio and remove physiological confounders, including detrending (linear, cubic and quadratic), regression of motion parameters and physiological noise including WM and CSF (29 regressors), despiking using 3DDESPIKE and low-pass filtering using a high-frequency cutoff of 0.15 Hz. We selected a 56s segmented window length based on previous work indicating that window lengths between 30–60 s successfully capture resting state dFC fluctuations and do not yield substantially different results, which, moreover, allowed us to match the window length of the LEMON dataset to the SALD dataset (i.e., 28*2=56 s) (Preti et al., 2017). We utilized the average sliding window correlation method for dFC (22s window length) to attenuate the impact of potential scanner drift, head motion, and physiological noise-related artifacts and to increase sensitivity to faster transient connectivity fluctuations, resulting in 284 (LEMON) and 198 (SALD) final correlation matrices containing 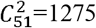 (LEMON) and 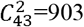 (SALD) component pairs (Vergara et al., 2019). A penalty on the L1 norm was imposed in the graphic LASSO framework with 100 repetitions to promote sparsity in estimation (Friedman et al., 2008). Sex was included as a covariate in the estimation. Finally, all resulting FC matrices were transformed to z-scores using Fisher’s z-transformation.

Next, we applied a K-means clustering algorithm to identify the number of distinct brain states (Lloyd, 1982). Time-series correlation matrices were used to estimate reoccurring FC. We used the L1 distance (Manhattan distance) function to estimate the similarity between windowed FC matrices (Aggarwal et al., 2001). The K-means algorithm was repeated 100 times to reduce the bias of initial random selection of cluster centroids (Kim et al., 2017). Moreover, windows consisting of local maxima in FC variance were used as subsamples for each participant to decrease redundancy and computational demands.

According to the previous literature on brain aging (Fiorenzato et al., 2019; Kim et al., 2017) and cluster number validity analysis of all subjects (Silhouette criterion (Rousseeuw, 1987)), the optimal number of clusters was decided to be two (k = 2). In line with K-means results, each participant’s dFC matrices were then categorized into one of the 2 clusters - or, states - based on their similarity to the cluster centroids. We also used Bayesian information (Warnick et al., 2018) and elbow criterion (Allen et al., 2014), two additional commonly-used clustering approaches, to validate the robustness of our results. These criteria revealed a slightly different number of distinct brain states (k=3: Bayesian information criteria; k=4: elbow criteria), although, importantly, all age-associated changes in temporal properties of dFC (fraction time, dwell time, discussed below) were observed when inspecting the results of alternative clustering methods (see Supplementary Results 2/Supplementary Table 1).

We extracted three temporal properties of each dFC state: 1) fraction time (i.e., the percentage of time spent in each state), 2) mean dwell time (state duration, i.e., the number of consecutive windows in a particular state before transferring another state), 3) and the number of transitions (i.e., how many times a switch between states occurred). The fraction, mean dwell time, and the number of transitions of the young and older groups were calculated using Wilcoxon rank-sum test.

#### 2.3.4 Dynamic topological metrics: global and local efficiency difference between two group

We used GRETNA (http://www.nitrc.org/projects/Gretna) to obtain topological properties of the resting state fMRI data. Brain aging is thought to affect two key graph metrics that control network efficiency: global and local efficiency (Kim et al., 2017; Xia et al., 2019). Global network efficiency was defined as the average efficiency across all node pairs, representing parallel information transfer in the network. In comparison, local efficiency was defined as the average of nodal local efficiency within neighbors of a node, which is thought to represent communication among neighbors.

LEMON and SALD ICs were defined as nodes and the FC between two components as edges for each time window. Subject-specific FC matrices were then binarized with respect to sparsity threshold (i.e., the number of actual edges divided by the maximum possible number of edges). The sparsity threshold was defined as 0.1 to 0.34 in 0.01 increments to maximize the reliability of the global and local efficiency estimation. In line with previous work (Kim et al., 2017), only included positive correlations in this study. Next, we obtained the global and local efficiency at each sparsity threshold. We applied an area under the curve (AUC) approach to avoid specific threshold selection (Kim et al., 2017; Navalpotro-Gomez et al., 2020; Xu et al., 2021). Finally, as suggested by Kim et al (Kim et al., 2017), we calculated the variance of the global and local efficiency over time to examine dynamic graph properties differences of younger and older participants and compared these metrics using Wilcoxon rank-sum test. For a visual representation of the data analysis pipeline, we refer the reader to Supplementary Figure 1.

## 3. Results

All results reported below were obtained using the entire sample of participants after the removal of exclusions according to the criteria described in section 2.1 (final sample: n LEMON=133, n SALD=330).

### 3.1 Sample descriptives

Demographics of each sample are reported in Table 1. We observed no significant sex differences between younger and older participants in either dataset, although older participants exhibited more head motion in both cohorts (Table 1 for all relevant statistics).

### 3.2 Brain network identity

#### 3.2.1 Brain network identity in the discovery cohort (LEMON)

The spatial maps of 51 independent components were identified using group ICA. We classified these components as belonging to one of seven networks: AU (auditory network: 2 ICs - component 13, 65), SM (sensorimotor network: 9 ICs - component 1, 2, 3, 4, 11, 16, 46, 64, 77), VI (visual network: 11 ICs - component 22, 25, 29, 33, 34, 38, 44, 69, 76, 84, 97), CEN (cognitive executive network: 18 ICs - component 17, 27, 28, 40, 42, 55, 57, 62, 67, 71, 73, 78, 79, 81, 86, 91, 95, 98), DM (default mode network: 4 ICs - component 18, 36, 51, 89), BG (basal ganglia network: 5 ICs - component 6, 7, 9, 20, 87), CB (cerebellar network: 2 ICs - component 50, 68). Spatial maps containing independent components and static between-network FC matrices are presented in Figure 1 (top row). More information regarding the location and statistics of independent components are presented in Supplementary Table 2.

**Figure 1 legend:**
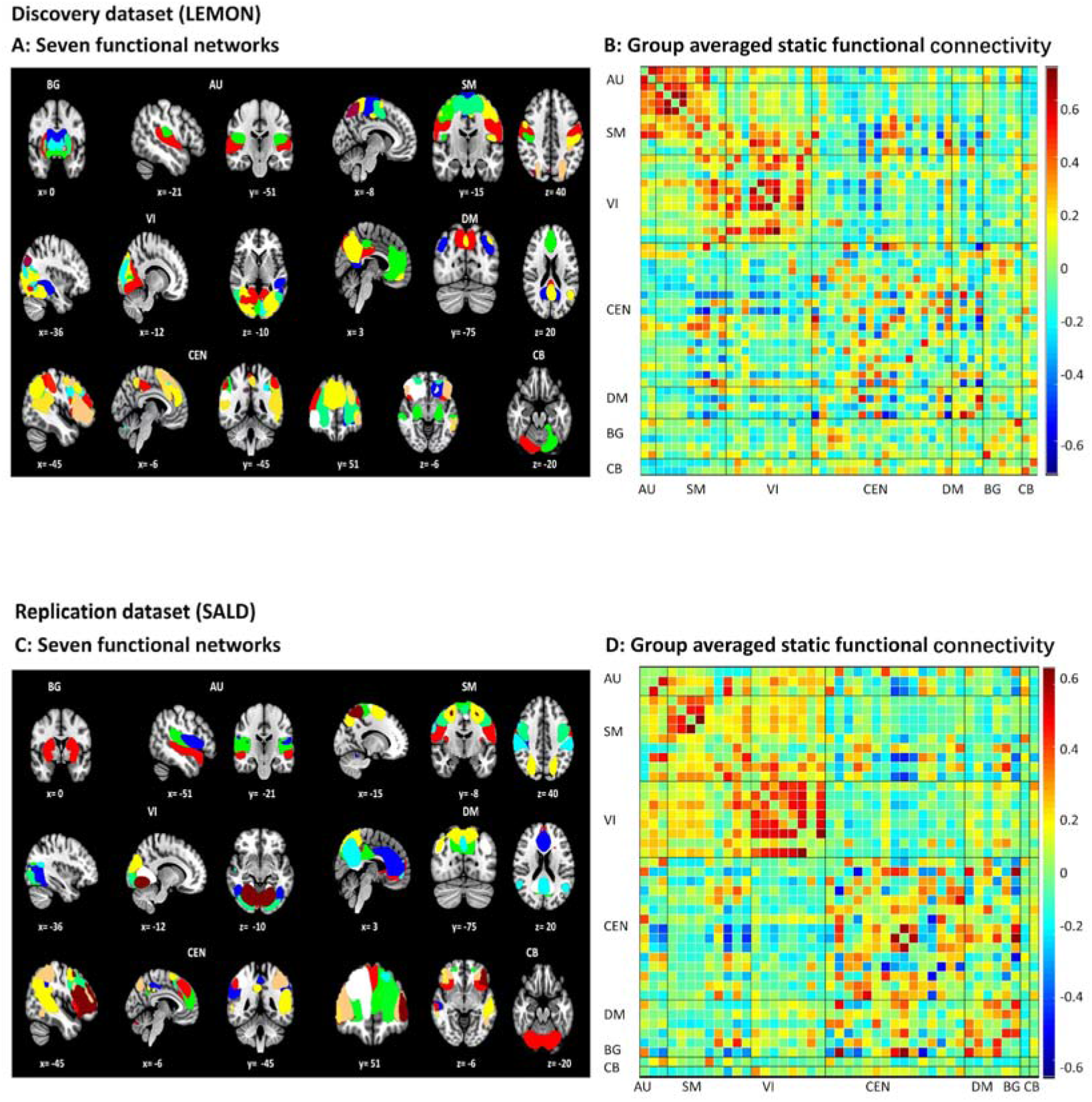
Independent components identified by group IC. A: LEMON/discovery cohort (top row) - 51 observed independent components across seven networks; B: static FC matrices for all 51 components; C: SALD/replication cohort (bottom row) - 43 observed independent components across seven networks; D: static FC matrices for all 43 components.

#### 3.2.2 Brain network identity in the replication cohort (SALD)

The spatial maps of 43 independent components in the SALD dataset were similarly assigned to one of seven networks: AU (auditory network: 3 ICs - component 58, 71, 98), SM (sensorimotor network: 9 ICs - component 3, 5, 11, 12, 44, 45, 64, 75, 80), VI (visual network: 8 ICs - component 17, 20, 29, 33, 42, 46, 62, 91), CEN: (cognitive executive network: 15 ICs - component 30, 34, 50, 55, 56, 59, 61, 67, 70,73, 76, 78, 79, 86, 92), DMN (default mode network: 6 ICs - component 8, 14, 21, 38, 54, 60), BG (basal ganglia network: 1 IC - component 19), CB (cerebellar network: 1 IC - component 13) (see Figure 1 bottom row for visualization and Supplementary Table 3 for location/statistics).

### 3.3 Age-dependent dFC changes are mostly replicable across cultures

#### 3.3.1 Temporal properties

##### Discovery cohort

Using k-means clustering, we uncovered two distinct functional brain states: 1) a more frequent and relatively sparsely connected state (state I; 74% of scan time), characterized by positive within-network correlations in SM, VI and DMN, and, 2) a less frequent between-network connected state (state II; 26% of scan time), characterized both by positive correlations between SM and VI, and negative correlations between SM and VI with DMN (Figure 2).

**Figure 2 legend:**
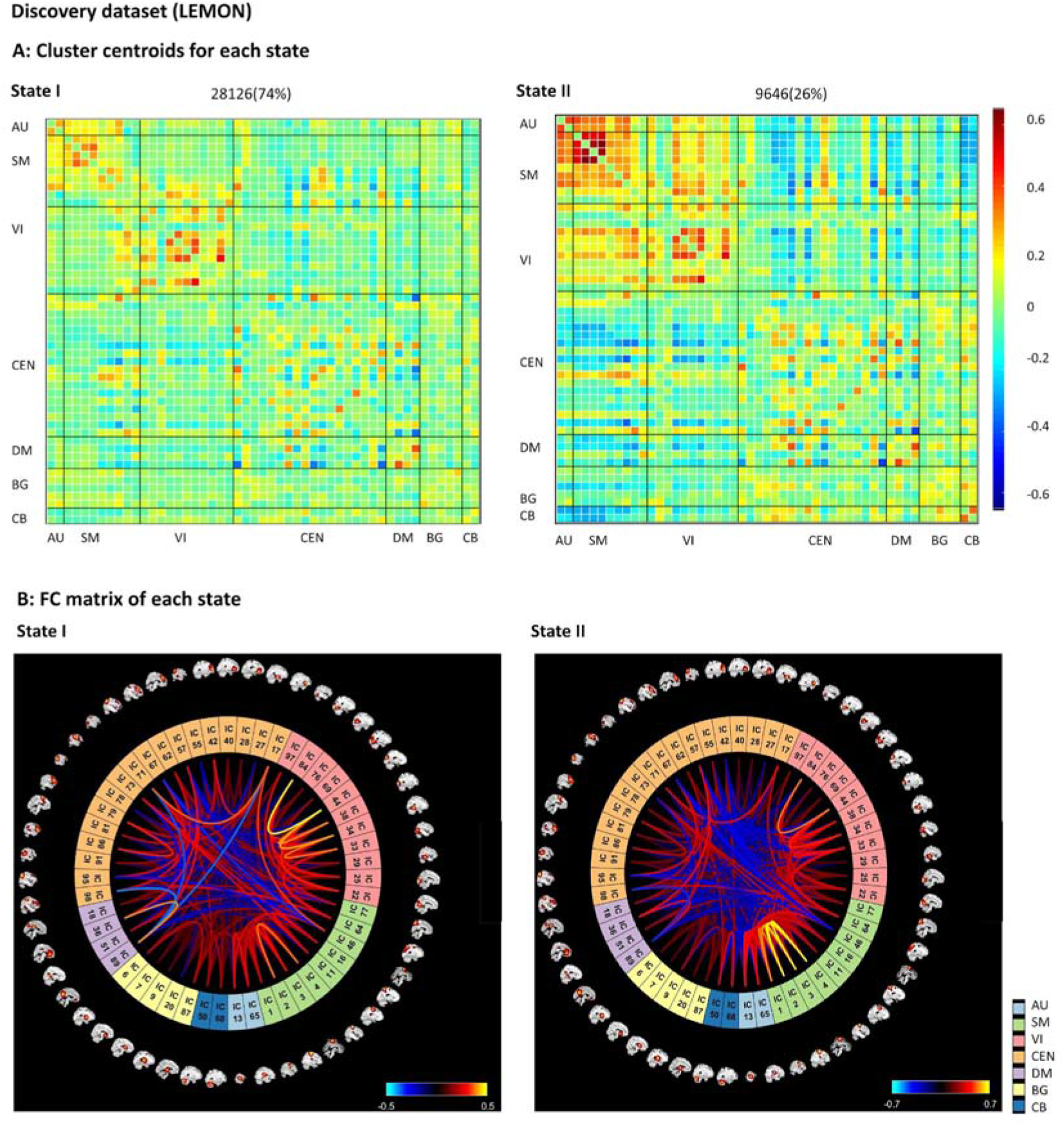
DFC brain states for LEMON dataset. A: The total number of occurrences and percentage of total occurrences are listed; B: FC matrix in each state.

Comparing younger and older individuals, we observed significant differences in fraction time for both states (state I: young: 79.19 ± 27.12%, older: 65.23 ±31.35%, z= 2.56, p=0.010, Cohen’s d= 0.46; state II: young: 20.81 ± 27.12%, older: 34.77 ± 31.35%, z= −2.56, p=0.010, Cohen’s d= −0.46, Wilcoxon rank-sum test), suggesting a 13.96% decrease/increase in state I/II in older participants. Older compared to younger participants additionally exhibited significantly shorter dwell time in state I (young: 164.19 ± 98.31, older: 116.12 ± 96.67, z= 2.82, p=0.004, Cohen’s d= 0.50, Wilcoxon rank-sum test), and higher dwell times in state II (young: 36.05±50.72, older: 53.60 ±59.63, z=-1.98, p=0.048, Cohen’s d= 0.35, Wilcoxon rank-sum test). Lastly, older compared to younger participants more frequently switched between distinct brain states (young: 1.86±1.85, older: 2.58 ±1.78, z=-2.55, p=0.011, Cohen’s d= 0.45, Wilcoxon rank-sum test), suggesting a reduction in the temporal stability of brain states in older participants (see Figure 4 top row).

##### Replication cohort

Similar to the LEMON dataset, we observed a more frequent (62%) relatively sparsely connected state characterized by a positive within-network correlation in SM, VI, CEN, and DMN (state I; 62% of scan time) and a less frequent but strongly between-network connected state (state II; 38% of scan time), characterized by positive correlations between SM and VI and negative correlations between SM and AU with DMN (Figure 3)

**Figure 3 legend:**
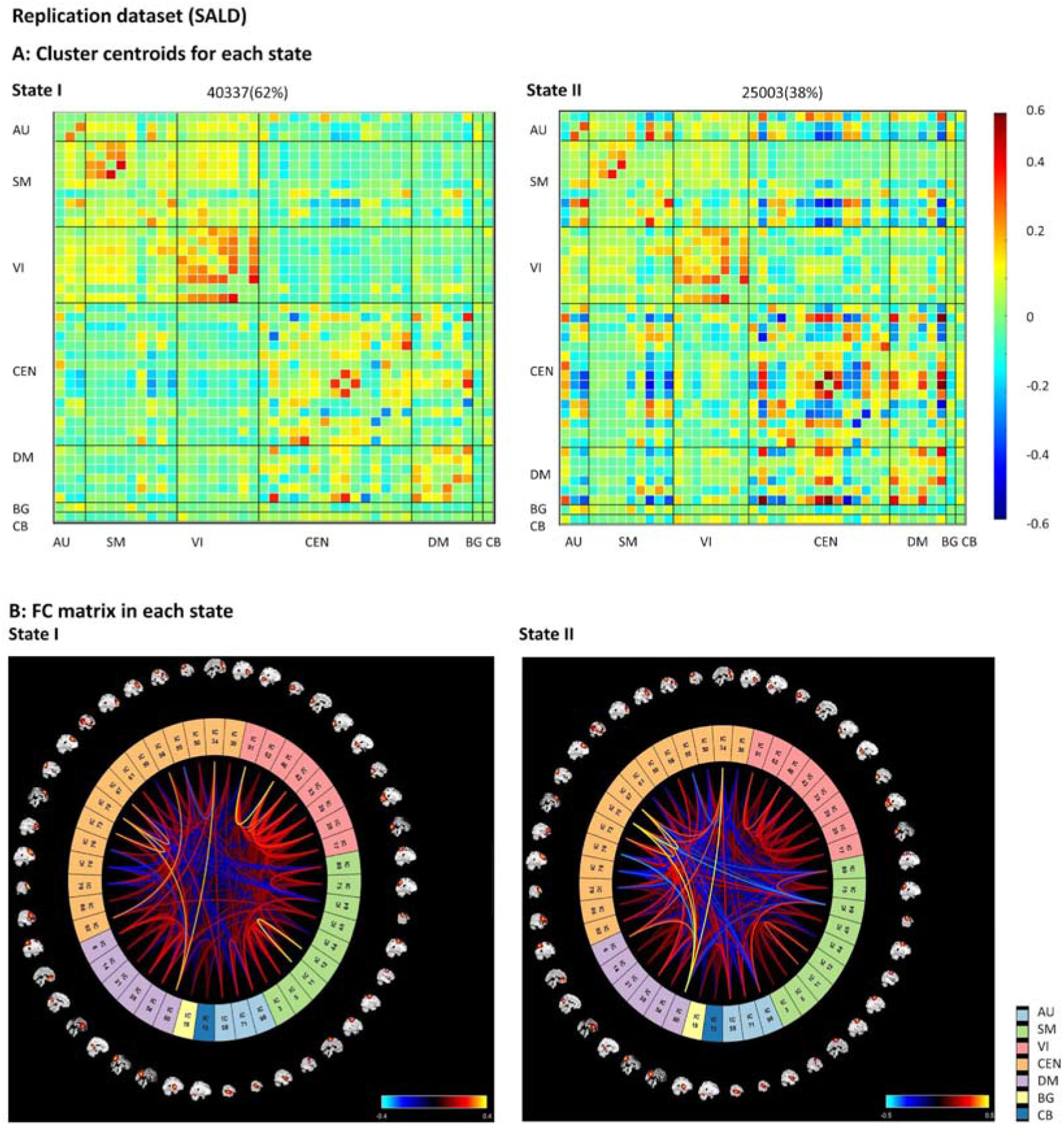
DFC brain states for SALD dataset. A: The total number of occurrences and percentage of total occurrences are listed; B: FC matrix in each state.

With respect to the key outcome dFC measures that differed between younger and older SALD participants, we replicated most observations in the LEMON dataset. That is, older participants spent less time in state I and more time in state II (state I: young: 65.75 ± 22.94’%, older: 57.57± 27.79%, z= 2.70, p=0.007, Cohen’s d= 0.30; state II: young: 34.25± 22.94%, older: 42.43±27.79%, z= −2.70, p=0.007, Cohen’s d= −0.30, Wilcoxon rank-sum test; 8.18 % decrease/increase in state I/II in the older participant). Older participants also had significantly shorted dwell times in state I (young: 65.23±52.42, older: 57.82±55.39, z=2.77, p=0.006, Cohen’s d=0.31, Wilcoxon rank-sum test), and a marginally longer dwell time in state II (young: 30.35±25.51, older: 38.43±38.43, z=-1.79, p=0.073, Cohen’s d=0.20). Older and younger participants, however, did not differ in number of state transitions (young: 3.83±2.21, older: 3.94 ±2.34, z=-0.63, p=0.528, Cohen’s d=0.07, Wilcoxon rank-sum test) although, numerically, the direction of brain state transitions in older individuals matched with observations from the LEMON dataset (see Figure 4 bottom row).

**Figure 4 legend:**
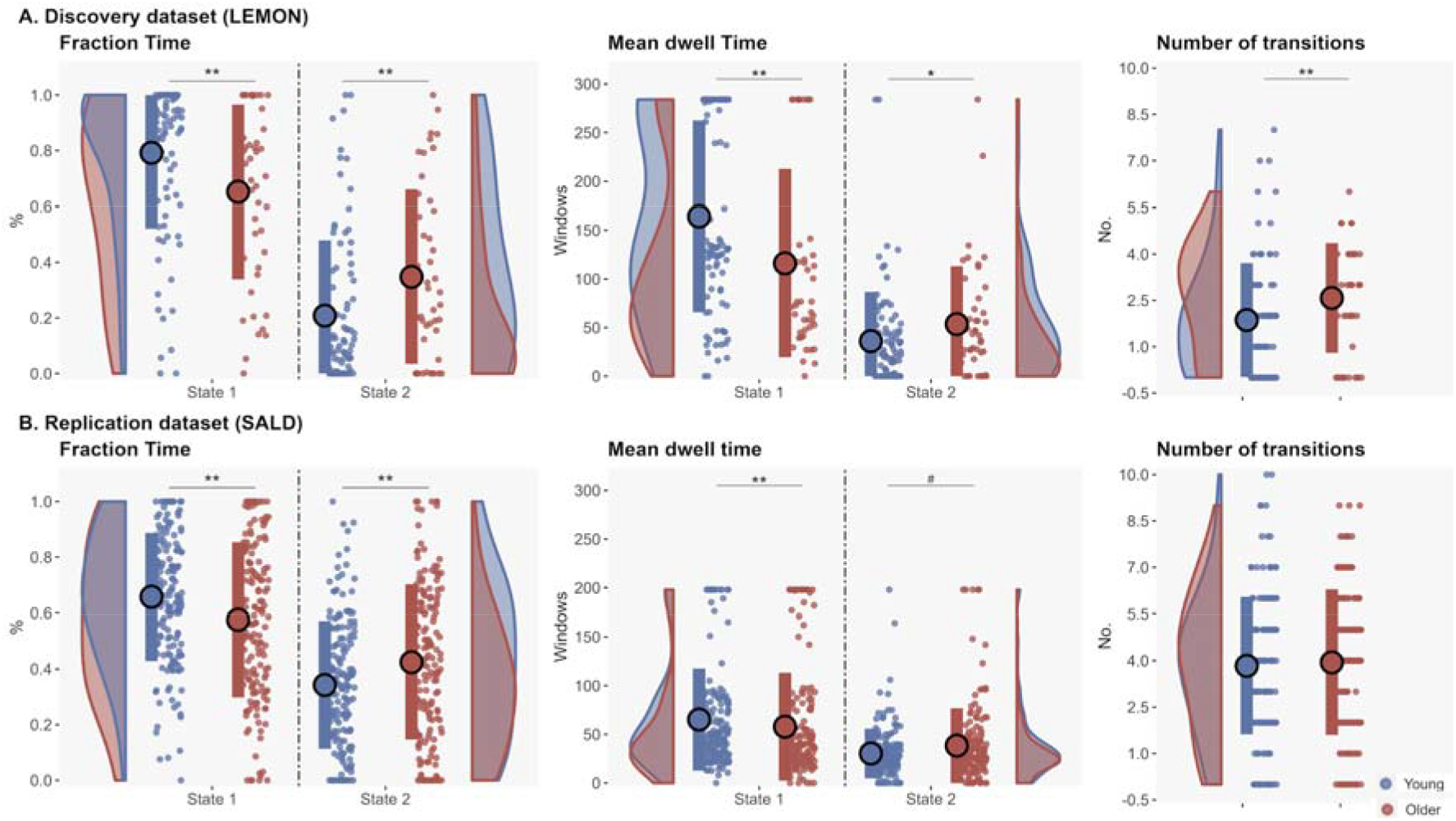
Age-dependent changes in the temporal properties of dFC (Fraction time; mean dwell time; number of transitions). A: LEMON dataset (top row); B: SALD (bottom row) dataset. (#: 0.05-0.1; *: p<0.05; **: p<0.01; ***: p< 0.001).

#### 3.3.2 Topological properties

##### Discovery cohort

We observed a significant difference in global efficiency variance between younger and older participants, with older participants showing higher global efficiency (young: 6.26×10^−5^ ± 7.47×10^−5^, older: 7.26×10^−5^ ± 4.88×10^−5^, z=-2.30, p=0.021, Cohen’s d =0.41, Wilcoxon rank-sum test). When excluding outliers with values ± 3SD above the group mean, the two groups still differed on global efficiency (young: 5.09×10^−5^ ± 3.85×10^−5^, older: 7.26×10^−5^ ± 4.88×10^−5^, z=-2.70, p=0.007, Cohen’s d=0.48, outliers=3). However, significant group differences in local efficiency were not observed (excluding ± 3SD outlier: young: 6.21×10^−5^ ± 2.95×10^−5^, older: 6.21×10^−5^ ± 2.66×10^−5^, z=-0.05, p=0.956, Cohen’s d=0.009; including ± 3SD outlier: young: 6.09×10-5 ± 2.74×10-5, older: 6.21×10-5 ± 2.66×10-5, z=-0.05, p=0.870, Cohen’s d=0.006, outliers=1) (see Figure 5 left row).

**Figure 5 legend:**
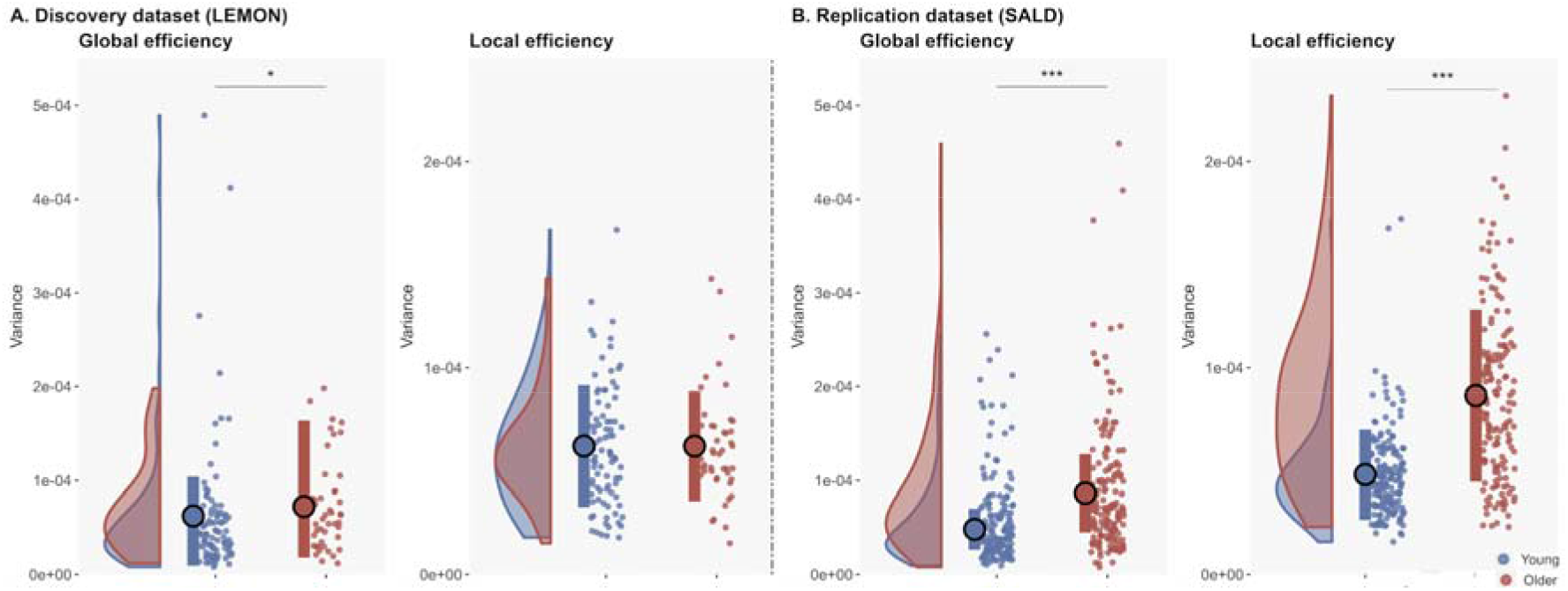
Age-dependent changes in temporal topological properties (variance of global and local efficiency). A: LEMON (left) dataset; B: SALD (right) dataset. (*: p<0.05; ***: p< 0.001).

##### Replication cohort

Older compared to younger participants exhibited higher global efficiency variance (young: 5.68× 10^−5^±4.74×10^−5^, older: 9.12×10^−5^±7.29×10^−5^, z=-5.90, p<0.001, Cohen’s d=0.69) and local efficiency (young: 4.83×10^−5^±2.18×10^−5^, older: 8.66×10^−5^±4.16×10^−5^, z=-9.39, p<0.001, Cohen’s d=1.29). These results were not affected by outliers with extreme (± 3SD) global/local efficiency values (global efficiency: young: 5.68×10^−5^±4.74×10^−5^, older: 8.28×10^−5^±5.46×10^−5^, z=-5.53, p<0.001, Cohen’s d=0.64, outliers=5; local efficiency: young: 4.83×10^−5^±2.18×10-5, older: 8.30×10-5±3.68×10-5, z=-9.11, p<0.001, Cohen’s d=1.16, outliers=5, Wilcoxon rank-sum test) (see Figure 5 right row). An overview of (non-)replicated results is provided in Table 2.

**Table 2 legend:** Overview of age-associated changes in dFC metrics in the discovery and replication cohort

| Finding | Replicated? |
| --- | --- |
| Fraction time |  |
| Aging-associated increases in fraction time in the between-network connectivity state. | Replicated |
| Mean dwell time |  |
| Aging-associated increases in mean dwell time in the between-network connectivity state. | Replicated |
| Number of transitions | Observed only in the |
| Aging-associated increases in brain state transitions | discovery cohort |
| Global efficiency |  |
| Aging-associated increases in global efficiency variance. | Replicated |
| Local efficiency | Observed only in the |
| Aging-associated increases in local efficiency variance. | replication cohort |

## 4. Discussion

We conducted the first cross-cultural replication study of age-dependent dFC changes in two independent samples, with the aim of assessing the robustness and reliability of dynamic functional brain changes associated with aging. In both the discovery and replication cohort, we found evidence for two distinct brain states: a dominant state characterized primarily by dense positive connectivity patterns within well-characterized functionally-distinct networks (e.g., CEN, DMN), as well as a less frequent pattern of high interconnectivity between these networks. Importantly, most temporal properties of these distinct connectivity states differed with age, such that younger individuals displayed higher fraction time and longer mean dwell time in the more frequent within-network connectivity state. In contrast, fraction and mean dwell time in the less functionally-specific between-network connectivity state were greater for older participants. Older individuals moreover tended to exhibit a greater number of transitions between these two distinct brain states, as well as greater variance in global efficiency. Together, these observations yield two important insights. Firstly, our results confirm that healthy aging is associated with a reorganization of dynamic brain activation, such that connectivity patterns that favor functional specificity are traded in for less efficient, whole-brain, connectivity patterns. Secondly, we demonstrate that these age-dependent dFC changes are relatively stable across samples with presumably different exposure to socio-cultural factors and health care access. Our results were also robust against variations in scan duration and clustering methods, suggesting that they were not the result of analytical idiosyncrasies.

Regarding the age-associated changes in temporal properties of connectivity states, we observed that older participants exhibited reduced reliance on segregated, functionally specialized, brain networks at rest. Numerous previous studies have reported that healthy older individuals exhibit alterations in FC; both within well-defined high-order cognitive networks (i.e., DMN, CEN), as well as primary processing networks (i.e. VI and SM) (Cassady et al., 2019; Chan et al., 2014; Damoiseaux, 2017; Hafkemeijer et al., 2012; Lim et al., 2014; Oschmann and Gawryluk, 2020; Tomasi and Volkow, 2012; Zonneveld et al., 2019). One way to interpret these aging-associated changes in network configurations is as a compensatory mechanism. For example, the STACr model (The Scaffolding Theory of Aging and Cognitive revised) posits that adverse effects on functional and structural brain processes can be ameliorated by ‘compensatory scaffolding’ (Reuter-Lorenz and Park, 2014). Our results fit well with this framework, which views reliance on between-network connectivity patterns as a compensatory mechanism that preserves cognitive functions at old age. Although speculative, aging-associated compensatory mechanisms in our sample could be reflected by increased/decreased fraction time and mean dwell time in the between/within-network connectivity state.

We additionally observed robust group differences in dynamic whole-brain topology measures that largely replicated across cultural backgrounds. Previous studies have demonstrated an overall decline in global and local efficiency, derived from static FC analyses, in older compared to younger adults (Gomez-Ramirez et al., 2015; Iordan et al., 2017; Varangis et al., 2019; Xia et al., 2019). However, to our knowledge, no previous studies have investigated age-associated variability in topological organization, obtained using dFC. Compared to their younger counterparts, older participants showed significantly higher global efficiency variability. Global efficiency metrics are thought to capture parallel information transfer and integrated processing, which correlate positively with IQ and cognitive abilities (Bullmore and Sporns, 2012). Thus, greater age-associated variability in global efficiency suggests that less time spent in brain states dominated by functionally-distinct networks results in more unstable, and thus less efficient, parallel information transfer (Bullmore and Sporns, 2012; Kim et al., 2017).

It has been hypothesized that culturally-specific factors (e.g., dietary habits, exposure to distinct risk factors, health care quality and accessibility) may influence the progression of neurodegenerative diseases and cognitive decline more generally (Rodriguez et al., 2022; Sheng et al., 2022). However, previous dFC studies have mainly been conducted on culturally homogenous samples (Tian et al., 2018; Viviano et al., 2017; Xia et al., 2019; Zhang et al., 2021b), raising doubts about the universality of age-related dFC changes. This first cross-culture dFC fMRI study shows broad similarities across cultures in terms of dFC temporal and topological properties of older participants. Specifically, our analyses reveal a consistent picture, independent of cultural background and analytical choices, of age-associated changes in temporal and topological dFC properties that hint at increased reliance on less functionally specialized and less efficient neural networks in healthy older participants. Taken together, this work suggests that dFC metrics, especially fraction time, dwell time and global efficiency variance, may be promising and generalizable neural indexes for brain changes in older healthy and pathological populations.

For example, Parkinson’s and associated diseases (i.e. Parkinson’s disease mild cognitive impairment – PDM; Parkinson’s disease dementia - PDD) are characterized by altered temporal properties in dynamic connectivity (Díez-Cirarda et al., 2018; Fiorenzato et al., 2019; Kim et al., 2017; Xu et al., 2021). Previous work found decreased mean dwell time in segregated within-network connectivity states and increased brain state transitions in PD and PDM compared to matched control participants (Díez-Cirarda et al., 2018; Kim et al., 2017). PDD, however, exhibited longer dwell time in segregated within-network connectivity states, in combination with fewer brain state transitions (Fiorenzato et al., 2019). These results underscore the potential of dFC’s sensitivity to distinct neurodegenerative states.

Despite the successful implementation of a novel cross-cultural neuroimaging replication approach, the present results need to be interpreted in the context of limitations. Firstly, the numbers of young and older participant groups in the LEMON dataset were unbalanced (higher number of young than older participants in the final analysis sample) due to use of more stringent exclusion criteria that are known to impact resting state and dFC metrics (e.g., motion). Secondly, resting state instructions differed between the two datasets: whereas in the LEMON dataset participants were instructed to keep their eyes open, participants in the SALD study were instructed to close their eyes. While several studies have revealed that these different data acquisition conditions impact FC and dFC metrics (Agcaoglu et al., 2019; Patriat et al., 2013; Weng et al., 2020), findings in the present study replicated successfully across these commonly used resting state conditions. Thirdly, the sliding window technique has limitations inherent to the methodology, such as the effect of window lengths on the results (Preti et al., 2017), which we aimed to minimize by relying on previously-suggested window lengths.

Taken together, this work provides novel insights into age-associated changes in dynamic neural processes across different cultural backgrounds, suggesting that both dynamic temporal properties (e.g., fraction time, dwell time) and topological (e.g., global efficiency) are robustly altered with advancing age, suggesting that such metrics may prove to be important functional correlates of brain aging.

## Supporting information

Age-dependent changes in the dynamic functional organization of the brain at rest_supplementary

## Acknowledgments

This work was supported by the China MOST2030 Brain Project (Grant No. 2022ZD0208500) and the China Scholarship Council (CSC) (Grant No. 202006070007).

## Author Contributions

X.Y. and X.Z. designed this study; X.Y. performed data analysis with the help of X.Z., D.H., B.B., D.L. and F.X.; interpretation of data and drafting of the manuscript was done by X.Y., D.H., B.B., D.L. All authors read and approved the final manuscript.

## Data and materials availability

All fMRI data included in this study are publicly available (LEMON dataset: http://fcon_1000.projects.nitrc.org/indi/retro/MPI_LEMON.html) and (SALD dataset: http://fcon_1000.projects.nitrc.org/indi/retro/sald.html). The dFC results of the current study are available in the Open Science Framework (https://osf.io/xenpc/).

