## Supplementary material for "Age-dependent changes in the dynamic functional organization of the brain at rest – a cross-cultural replication approach": Age-dependent changes in the dynamic functional organization of the brain at rest_supplementary

**Supplementary materials**

**Supplementary results**

**1. Results - brain states obtained using the entire timeseries (LEMON dataset)**

When using the entire resting state fMRI time series, we obtained highly similar distinct brain states. Compared with the analyses reported in the main text, we observed a slight difference in relative time spent in each state. Specifically, we observed (1) a more frequent and relatively sparsely connected state (state I; 78% of scan time), characterized by positive within-network correlations in SM, VI and DMN and (2) a less frequent state with higher between-network connectivity (state II; 22% of scan time), characterized both by positive correlations between SM and VI, and negative correlations between SM and VI with DMN (see Supplementary Figure 2).

**2. Results - group comparisons of temporal properties using different cluster solutions (K=3 and K=4) in both cohorts**

When following up the results from the cluster estimation algorithms that yielded a different number of distinct brain states (k=3 or k=4), we still observed a frequently connected within-network (State I) and a less frequent but interconnected between-network state (state II) in both cohorts. Under any cluster solutions (K=2 or K=3 or K=4), we found that in the LEMON dataset, state I presented as a within-network FC configuration, particularly located in the SMN, VI, and DMN networks, and state II was characterized by stronger between-network FC configurations that included positive correlations between the SMN and VI networks, as well as anti-correlations between the SMN and VI with DMN networks (Supplementary Figure 3 and Figure 4 top row). In the SALD dataset, state I was characterized by positive couplings located mainly within networks (i.e. SMN, DMN, CEN and VI networks), state II was characterized mainly by stronger positive couplings between networks (i.e. SMN and VI), as well as anti-related correlations (i.e. SM and DMN & and AU and DMN) (Supplementary Figure 3 and Figure 4 bottom row). Thus, these cluster estimation methods yielded two states (State I and II for each solution) that resembled those reported in the main text.

Moreover, similar to the results reported in the main text, we observed reduced/increased fraction time and mean dwell time (State I/II) in older compared to younger individuals. In contrast to the main text result, we observed no age differences in the number of brain state transitions (Supplementary Table 1).

**Supplementary figures**

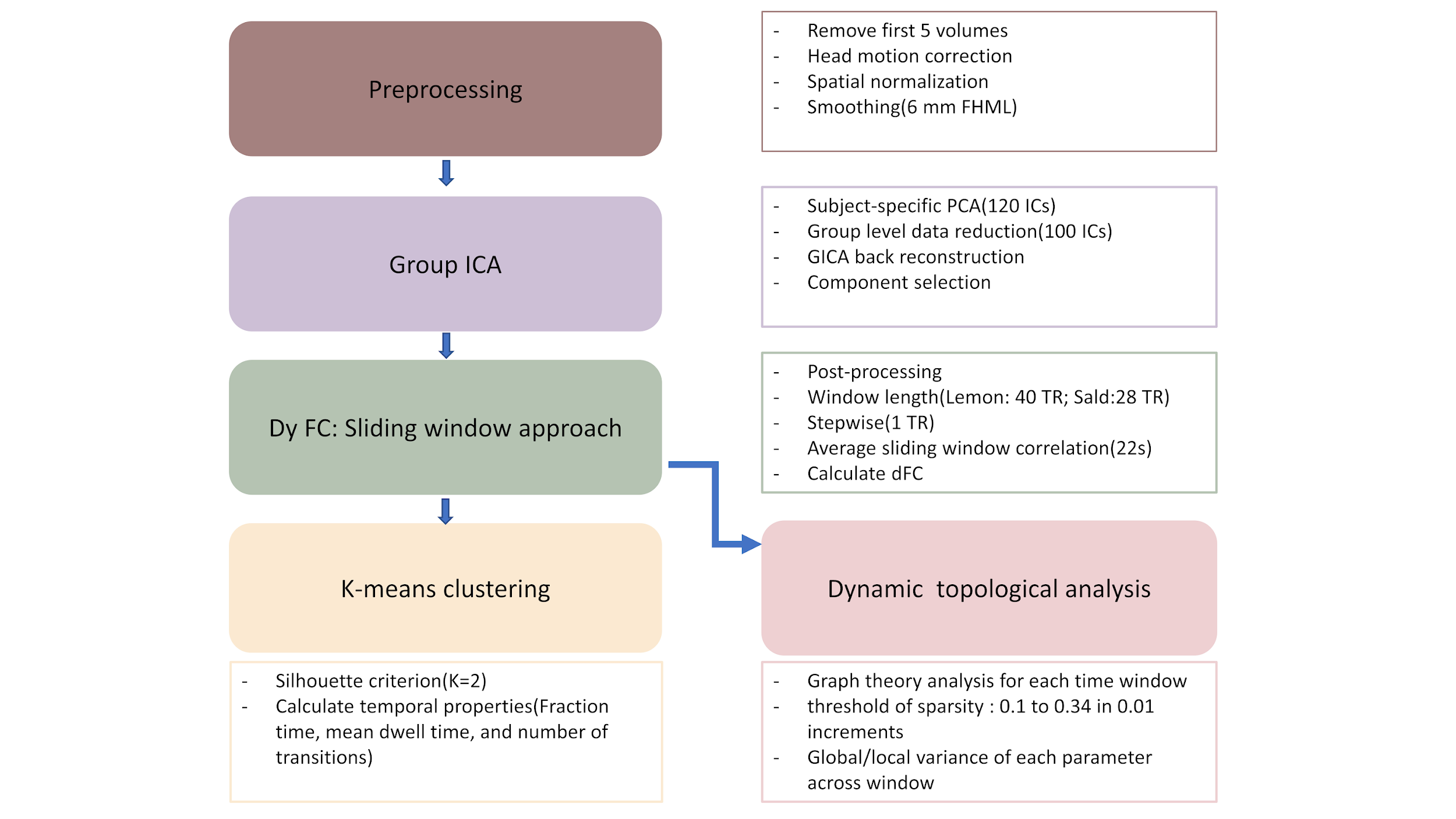

Supplementary Figure 1. Data analysis pipeline

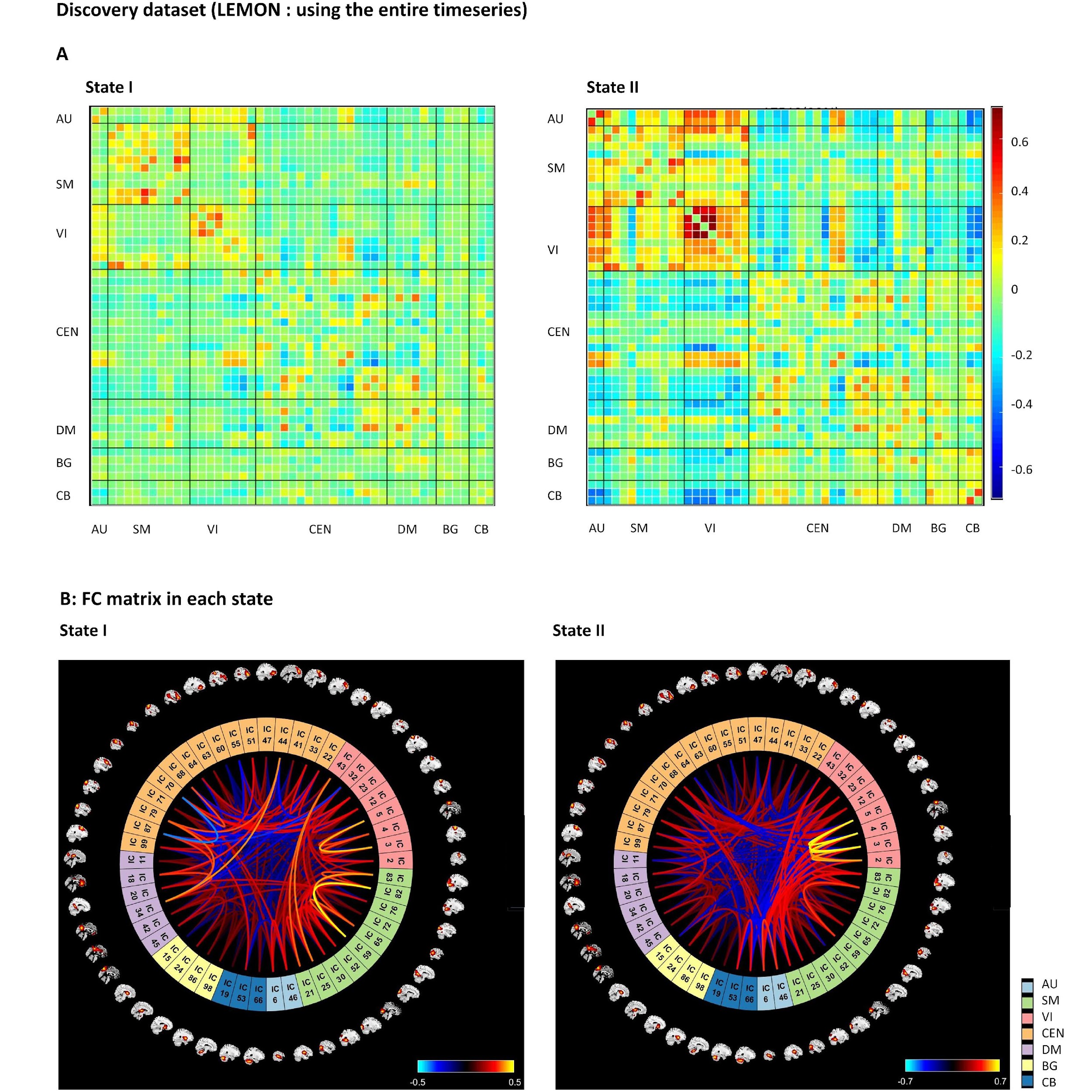

Supplementary Figure 2 Legend: DFC brain states obtained using the entire timeseries (LEMON dataset). A: The total number of occurrences and percentage of total occurrences are listed; B: FC matrix in each state.

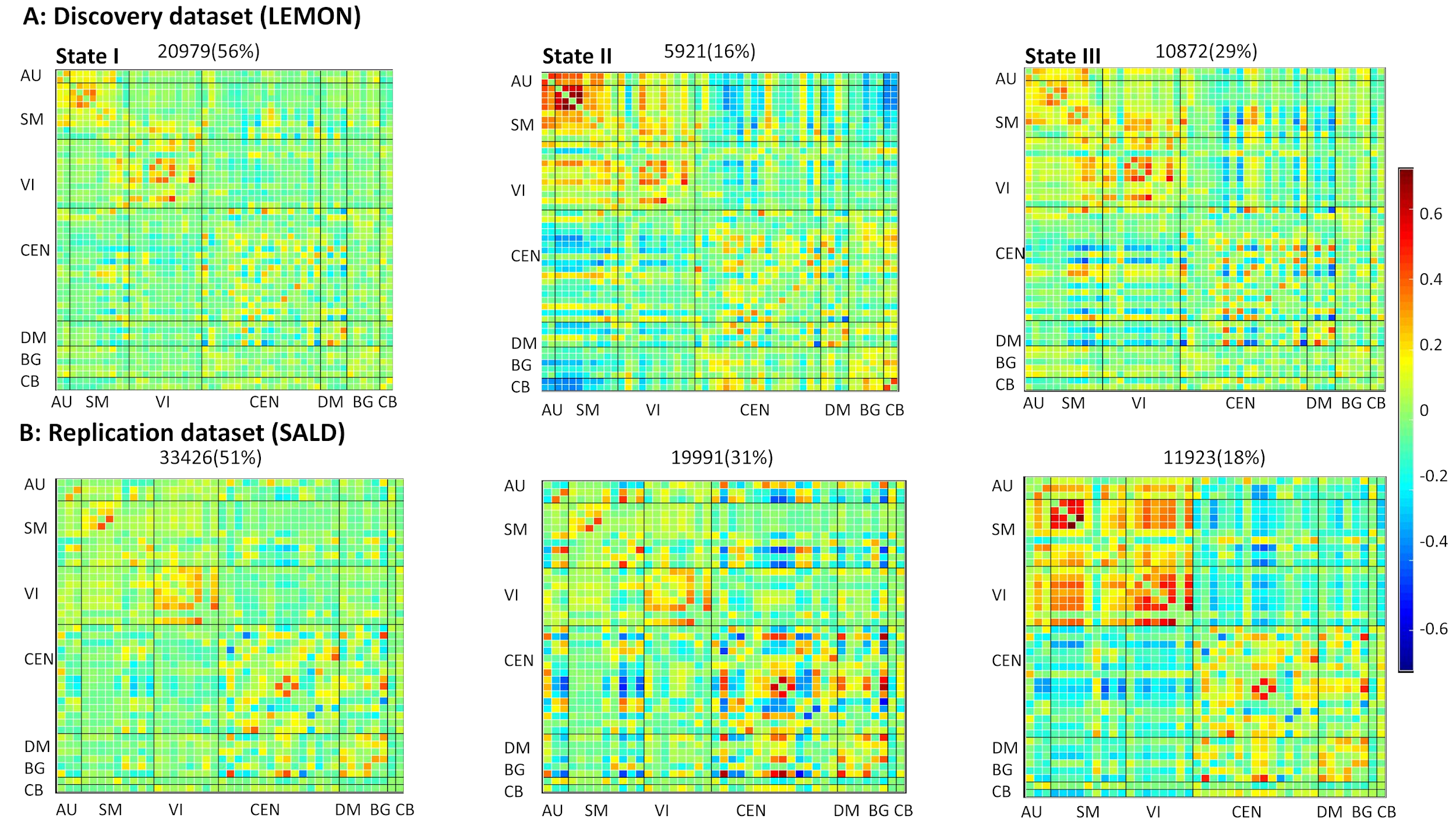

Supplementary Figure 3 Legend: Results of 3-cluster solution. The total number of occurrences and percentage of total occurrences are listed above each cluster median. (A: LEMON dataset - top row; B: SALD dataset - bottom row).

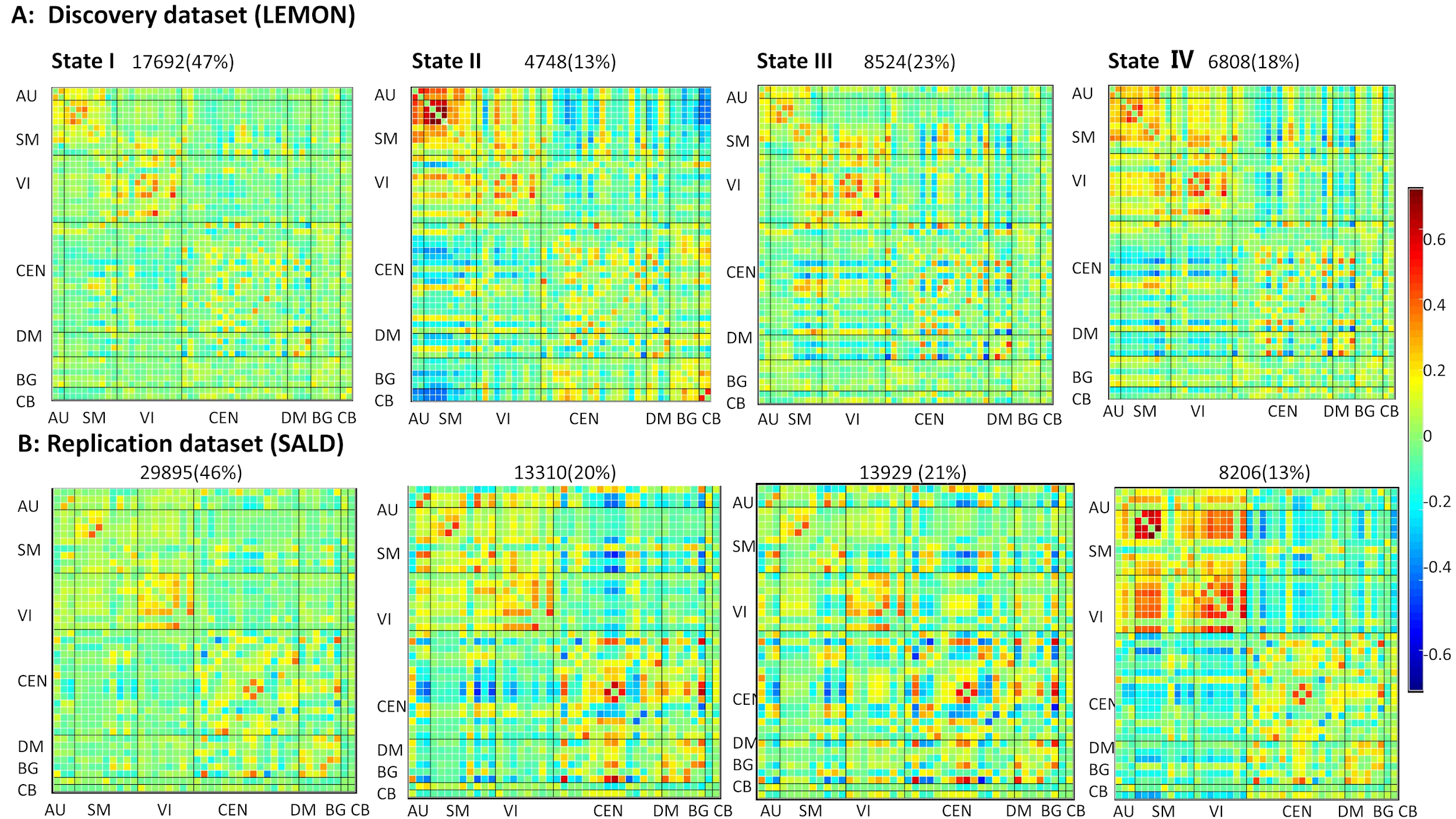

Supplementary Figure 4 Legend: Results of 4-cluster solution. The total number of occurrences and percentage of total occurrences are listed above each cluster median. (A: LEMON dataset - top row; B: SALD dataset - bottom row).

**Supplementary tables**

| **K=3**  LEMON  SALD |  | Young | Older | T  df=131  df=328 | P | Cohen's d |
| --- | --- | --- | --- | --- | --- | --- |
| Fraction | state 1 | 58.46±26.98 | 49.84±30.24 | 1.67 | 0.096 # | 0.31 |
|  |  | 55.62±24.73 | 46.53±28.23 | 3.11 | 0.002 ** | 0.34 |
|  | state 2 | 11.76±20.76 | 23.34±33.57 | -2.45 | 0.015 * | 0.45 |
|  |  | 27.18±21.01 | 34.13±26.17 | -2.67 | 0.008 ** | 0.29 |
|  | state 3 | 29.79±23.59 | 26.82±25.08 | 0.67 | 0.504 | 0.12 |
|  |  | 17.20±21.34 | 19.33±24.15 | -0.85 | 0.400 | 0.09 |
| Dwell Time | state 1 | 75.03±59.76 | 74.57±73.14 | 0.04 | 0.969 | 0.01 |
|  |  | 45.86±41.98 | 40.02±42.50 | 1.25 | 0.209 | 0.14 |
|  | state 2 | 17.50±30.41 | 40.08±62.52 | -2.81 | 0.006 ** | 0.52 |
|  |  | 24.37±22.16 | 30.32±30.84 | -2.02 | 0.044 * | 0.22 |
|  | state 3 | 37.32±28.15 | 39.07±52.16 | -0.25 | 0.801 | 0.05 |
|  |  | 18.65±24.05 | 19.99±26.76 | -0.48 | 0.633 | 0.05 |
| Transition |  | 4.34±2.01 | 3.82±2.14 | 1.38 | 0.170 | 0.25 |
|  |  | 5.29±2.42 | 5.07±2.44 | 0.81 | 0.418 | 0.09 |
| **K=4**  LEMON  SALD |  |  |  |  |  |  |
| Fraction | state 1 | 50.42±27.37 | 39.83±28.45 | 2.08 | 0.039 * | 0.38 |
|  |  | 49.53±24.26 | 41.48±27.15 | 2.72 | 0.007 ** | 0.30 |
|  | state 2 | 8.25±18.11 | 21.01±32.31 | -2.92 | 0.004 ** | 0.53 |
|  |  | 17.85±18.06 | 22.99±20.81 | -2.40 | 0.017* | 0.26 |
|  | state 3 | 23.22±19.08 | 21.29±17.73 | 0.56 | 0.573 | 0.10 |
|  |  | 21.18±18.49 | 21.46±20.23 | -0.14 | 0.892 | 0.02 |
|  | state 4 | 18.10±19.14 | 17.87±20.52 | 0.06 | 0.949 | 0.01 |
|  |  | 11.44±18.74 | 13.71±22.01 | -1.00 | 0.313 | 0.11 |
| Dwell time | state 1 | 64.81±62.00 | 55.94±63.94 | 0.77 | 0.441 | 0.14 |
|  |  | 40.19±33.69 | 36.43±40.14 | 0.92 | 0.356 | 0.10 |
|  | state 2 | 11.97±26.02 | 32.90±58.08 | -2.87 | 0.005 ** | 0.53 |
|  |  | 17.17±18.49 | 21.15±20.83 | -1.84 | 0.067# | 0.20 |
|  | state 3 | 30.57±21.66 | 28.24±22.97 | 0.58 | 0.566 | 0.11 |
|  |  | 21.01±18.39 | 19.93±18.64 | -0.14 | 0.892 | 0.02 |
|  | state 4 | 26.40±30.96 | 28.12±40.17 | -0.27 | 0.785 | 0.05 |
|  |  | 12.97±20.29 | 15.84±29.91 | -0.14 | 0.598 | 0.02 |
| Transitions |  | 5.51±2.44 | 5.33±2.84 | 0.38 | 0.707 | 0.07 |
|  |  | 6.05±2.53 | 6.20±2.76 | -0.54 | 0.592 | 0.06 |

Table 1 legend: Results of temporal properties when K (number of cluster is 3 and 4) (mean ± SD).

(#: p=0.05-0.09; *: p<0.05; **: p<0.01; ***: p<0.001; two-sample T test).

**Supplementary Table 2**

| **AU (2)** |  | **BA** | **K** | **T-value** | **MNI** |
| --- | --- | --- | --- | --- | --- |
| IC 13 | R superior temporal gyrus | 42 | 1863 | 46.00 | 63 -24 9 |
|  | L superior temporal gyrus | 41 | 1701 | 39.63 | 48 -27 9 |
| IC 65 | L postcentral gyrus | 43 | 1366 | 27.39 | -39 -18 18 |
|  | L postcentral gyrus | 43 | 1742 | 27.25 | 39 -18 18 |
| **SM (9)** | |  |  |  |  |
| IC 01 | L precentral gyrus | 6 | 1128 | 83.58 | -54 -9 33 |
|  | R precentral gyrus | 6 | 1208 | 78.09 | 57 -6 30 |
| IC 02 | R precentral gyrus | 4 | 2511 | 54.64 | 39 -18 57 |
| IC 03 | Medial frontal gyrus | 6 | 2193 | 58.84 | 3 -27 66 |
| IC 04 | L postcentral gyrus | 3 | 2230 | 57.51 | -42 -27 57 |
| IC 11 | Medial frontal gyrus | 6 | 2914 | 32.27 | 24 -9 63 |
| IC 16 | B postcentral gyrus | 5 | 3649 | 42.87 | 21 -45 66 |
| IC 46 | Superior frontal gyrus | 6 | 3038 | 35.88 | 6 3 63 |
|  | L precentral gyrus | 6 | 1556 | 30.90 | -45 -9 51 |
| IC 64 | B superior parietal lobule | 7 | 5517 | 37.98 | -21 -63 60 |
| IC 77 | M precuneus | 7 | 7192 | 46.09 | 6 -60 60 |
| **VI (11)** | |  |  |  |  |
| IC 22 | B calcarine_ | 31 | 3917 | 65.06 | 9 -69 15 |
| IC 25 | B cuneus | 17 | 2789 | 43.90 | -3 -87 3 |
| IC 29. | L middle temporal gyrus | 21 | 1920 | 32.29 | -45 -42 -12 |
|  | R temporal lobe | 21 | 1054 | 17.94 | 51 -42 -12 |
| IC 33 | R middle temporal gyrus | 37 | 1927 | 38.69 | 48 -66 3 |
|  | L middle occipital gyrus | 19 | 1665 | 34.33 | -45 -72 3 |
| IC 34 | B fusiform | 36 | 4444 | 36.09 | 24 -42 -12 |
| IC 38 | B middle occipital gyrus | 18 | 2765 | 51.01 | -30 -90 -3 |
| IC 44 | B cuneus | 19 | 3521 | 45.77 | 21 -87 24 |
| IC 69 | L fusiform | 18 | 3741 | 26.65 | -27 -72 -9 |
| IC 76 | R calcarine | 30 | 2133 | 28.18 | 30 -69 9 |
|  | Posterior Cingulate | 30 | 1515 | 24.47 | -24 -72 9 |
| IC 84 | R fusiform | 18 | 3050 | 37.96 | 27 -69 -6 |
| IC 97 | R parietal lobe | 39 | 2587 | 24.20 | 36 -69 33 |
|  | Parietal lobe | 19 | 1032 | 22.33 | -48 -60 -6 |
| **CEN (18)** | |  |  |  |  |
| IC 17 | R insula | 45 | 1192 | 44.12 | 33 24 3 |
|  | L insula | 45 | 1027 | 37.63 | -30 24 0 |
|  | Cingulate Gyrus | 32 | 1464 | 25.37 | 3 21 45 |
| IC 27 | B hippocampus | 28 | 3223 | 35.89 | -27 -15 -12 |
| IC 28 | L frontal Lobe | 11 | 2661 | 32.41 | -18 36 -6 |
| IC 40 | R frontal Lobe | 11 | 3319 | 32.66 | 18 33 -3 |
| IC 42 | B middle frontal gyrus | 10 | 4417 | 35.12 | -33 54 12 |
| IC 55 | R supramarginal gyrus | 40 | 1568 | 38.93 | 57 -45 36 |
|  | L supramarginal gyrus | 40 | 1717 | 35.75 | -54 -51 36 |
| IC 57 | Medial frontal gyrus | 9 | 4543 | 28.64 | 3 45 33 |
| IC 62 | L middle frontal gyrus | 46 | 2693 | 35.53 | -48 18 30 |
|  | R middle frontal gyrus | 46 | 1101 | 24.32 | 48 27 24 |
| IC 67 | L superior parietal lobule | 7 | 2915 | 36.05 | -36 -72 45 |
|  | L superior frontal gyrus | 8 | 2648 | 33.60 | -24 24 54 |
| IC 71 | M cingulate | 31 | 4498 | 27.96 | 3 -33 48 |
|  | L middle frontal gyrus | 46 | 1109 | 24.70 | -45 42 18 |
| IC 73 | L postcentral gyrus | 40 | 3281 | 42.54 | -51 -27 42 |
|  | R postcentral gyrus | 2 | 2201 | 34.54 | 60 -21 33 |
| IC 78 | R superior temporal gyrus | 13 | 3630 | 33.21 | 60 -45 18 |
| IC 79 | R inferior frontal gyrus | 46 | 2672 | 31.94 | 51 27 18 |
| IC 81 | M superior frontal gyrus | 6 | 3522 | 37.76 | -6 15 63 |
| IC 86 | L inferior frontal gyrus | 45 | 2334 | 40.88 | -54 24 15 |
|  | Inferior frontal gyrus | 47 | 748 | 17.31 | 51 36 -3 |
| IC 91 | L superior temporal gyrus | 22 | 2827 | 28.57 | -57 -48 9 |
|  | R superior temporal gyrus | 22 | 1211 | 20.76 | 60 -36 15 |
| IC 95 | R angular | 40 | 2431 | 47.52 | 45 -63 36 |
|  | R superior frontal gyrus | 8 | 2370 | 33.50 | 21 30 48 |
| IC 98 | Inferior parietal lobule | 40 | 2337 | 43.45 | 39 -39 45 |
|  | Middle frontal gyrus | 46 | 2457 | 27.35 | 45 39 18 |
| **DM (4)** |  |  |  |  |  |
| IC 18 | B precuneus | 7 | 5032 | 57.44 | 12 -66 36 |
| IC 36 | M anterior cingulate | 32 | 4201 | 41.77 | -3 42 -6 |
| IC 51 | B posterior cingulate | 29 | 3621 | 51.76 | -15 -57 18 |
| IC 89 | M precuneus | 7 | 3310 | 50.03 | 0 -63 33 |
| **BG (5)** |  |  |  |  |  |
| IC06 | R putamen | / | 1054 | 58.62 | 24 9 -3 |
|  | L putamen | / | 1024 | 56.08 | -24 6 3 |
| IC 07 | B putamen | / | 2125 | 52.13 | -15 9 -6 |
| IC 09 | B caudate | / | 2365 | 53.70 | 12 3 12 |
| IC 20 | L putamen | / | 1811 | 31.24 | -27 -12 6 |
| IC 87 | Medial dorsal nucleus | / | 2331 | 34.45 | -6 -15 9 |
| **CB (2)** |  |  |  |  |  |
| IC 50 | B cerebellum | / | 2986 | 50.84 | 30 -72 -33 |
| IC 68 | L cerebellum | / | 2336 | 31.21 | -30 -75 -39 |

Table 2 legend: MNI coordinates and statistics for LEMON dataset. BA = Brodmann area; L = left; R= right; M = medial; B = bilateral; K = cluster size.

**Supplementary Table 3**

| **AU (3)** |  | BA | K | T-value | MIN |
| --- | --- | --- | --- | --- | --- |
| IC 58 | R Superior Temporal Gyrus | 21 | 2752 | 45.69 | 60 -21 0 |
|  | L Middle Temporal Gyrus | 22 | 2938 | 36.31 | -60 -12 3 |
| IC 71 | R Postcentral Gyrus | 40 | 3318 | 50.13 | 60 -24 15 |
|  | L Temporal Gyrus | 41 | 3310 | 43.54 | -51 -27 12 |
| IC 98 | R Precentral Gyrus | 13 | 4450 | 40.47 | 42 0 9 |
|  | L Precentral Gyrus | 44 | 3313 | 39.06 | -45 0 6 |
| **SM (9)** |  |  |  |  |  |
| IC 3 | L Precentral Gyrus | 6 | 1664 | 82.68 | -51 -9 33 |
|  | R Precentral Gyrus | 6 | 1980 | 77.51 | 51 -6 33 |
| IC 5 | Medial Frontal Gyrus | 6 | 3308 | 62.13 | 0 -33 60 |
| IC 11 | R Precentral Gyrus | 4 | 3466 | 68.90 | 39 -21 63 |
| IC 12 | L Postcentral Gyrus | 3 | 3869 | 64.26 | -39 -24 57 |
| IC 44 | B Precentral Gyrus | 6 | 8994 | 46.61 | -45 -9 51 |
| IC 45 | B Middle Frontal Gyrus | 6 | 10155 | 57.33 | -24 0 7 |
| IC 64 | R Postcentral Gyrus | 2 | 5457 | 53.02 | 63 -21 33 |
|  | L Inferior Parietal Lobule | 40 | 5256 | 44.48 | -60 -24 30 |
| IC 75 | B Superior Parietal Lobule | 7 | 9708 | 48.61 | -21 -69 45 |
| IC 80 | B Postcentral Gyrus | 7 | 8054 | 55.83 | -21 -51 66 |
| **VI (8)** |  |  |  |  |  |
| IC 17 | R Calcarine | 17 | 5086 | 71.78 | 6 -87 3 |
| IC 20 | R Middle Occipital Gyrus | 18 | 7098 | 36.81 | -30 -84 6 |
| IC 29 | R Middle Temporal Gyrus | 37 | 2960 | 45.84 | 51 -66 0 |
|  | L Middle Occipital Gyrus | 37 | 2687 | 40.41 | -48 -72 0 |
| IC 33 | B Calcarine | 30 | 6827 | 50.88 | -18 -54 3 |
| IC 42 | B Middle Occipital Gyrus | 18 | 4936 | 62.88 | 30 -87 0 |
| IC 46 | L Cuneus | 19 | 6367 | 59.25 | -9 -90 27 |
| IC 62 | R Superior Temporal Gyrus | 13 | 4435 | 50.58 | 57 -45 15 |
|  | L Superior Temporal Gyrus | 22 | 5110 | 27.36 | -54 -51 9 |
| IC 91 | B Lingual Gyrus | 18 | 5974 | 52.60 | 21 -69 6 |
| **CEN (15)** |  |  |  |  |  |
| IC 30 | B Inferior Frontal Gyrus | 47 | 6429 | 52.35 | -30 27 3 |
| IC 34 | Medial Frontal Gyrus | 9 | 4951 | 60.21 | -3 54 21 |
| IC 50 | B Inferior Parietal Lobule | 40 | 13912 | 53.34 | -57 -48 39 |
| IC 55 | R Inferior Parietal Lobule | 40 | 6935 | 61.58 | 45 -42 51 |
|  | L Inferior Parietal Lobule | 40 | 3544 | 28.57 | -48 -45 51 |
| IC 56 | L Middle Frontal Gyrus | 46 | 11502 | 56.56 | -42 18 30 |
| IC 59 | B Middle Frontal Gyrus | 10 | 4046 | 47.37 | 30 60 12 |
| IC 61 | L Superior Temporal Gyrus | 22 | 5846 | 49.31 | -57 42 6 |
| IC 67 | B Middle Frontal Gyrus | 8 | 10125 | 54.27 | -24 30 45 |
| IC 70 | B Inferior Parietal Lobule | 40 | 14446 | 63.22 | 48 -63 39 |
| IC 73 | Medial Frontal Gyrus | 6 | 7910 | 50.61 | 3 36 42 |
| IC 76 | B Middle Frontal Gyrus | 43 | 10950 | 43.67 | 45 45 12 |
| IC 78 | R Middle Frontal Gyrus | 8 | 5761 | 34.65 | 24 30 45 |
| IC 79 | L Inferior Frontal Gyrus | 44 | 3022 | 44.72 | -54 15 21 |
|  | R Inferior Frontal Gyrus | 45 | 3180 | 28.97 | 57 21 21 |
| IC 86 | R Inferior Frontal Gyrus | 45 | 7561 | 50.48 | 48 24 27 |
| LC 92 | L Superior Parietal Lobule | 7 | 14944 | 46.55 | -36 60 54 |
| **DMN (6)** |  |  |  |  |  |
| IC 8 | Anterior Cingulate | 32 | 3536 | 61.24 | -3 39 -6 |
| IC 14 | Precuneus | 7 | 6489 | 80.38 | 12 -66 33 |
| IC 21 | Anterior Cingulate | 24 | 3527 | 56.26 | -3 33 6 |
| IC 38 | Precuneus | 23 | 7794 | 75.60 | 15 -54 18 |
| IC 54 | Precuneus | 7 | 7347 | 69.78 | 0 -60 54 |
| IC 60 | Precuneus | 7 | 8853 | 73.00 | 0 -60 33 |
| **SC (1)** |  |  |  |  |  |
| IC 19 | B Nucleus | / | 3242 | 63.54 | -27 0 3 |
| **CB (1)**  IC 13 | Cerebellum | / | 4196 | 55.36 | 24 -72 -24 |

Table 3 legend: MNI coordinates and statistics for SALD dataset. BA = Brodmann area; L = left; R= right; M = medial; B = bilateral; K = cluster size.
